# Superoxide dismutase impacts extracellular vesicle biogenesis and uptake

**DOI:** 10.1101/2025.11.04.686557

**Authors:** Nahin Siara Prova, Malek Elsayyid, Jessica E. Tanis

## Abstract

Extracellular vesicles (EVs), which transfer bioactive macromolecules between cells, play an important role in the pathogenesis of multiple neurodegenerative diseases. Focus has centered on how altered EV contents propagate disease and the potential for EVs as diagnostic biomarkers, while the effect of pathogenic factors on EV release is less understood. Here, we defined how the key antioxidant enzyme superoxide dismutase 1 (SOD-1) affects EV shedding from sensory neuron primary cilia, enrichment of ciliary proteins packaged into EVs, and uptake of EVs by surrounding glia *in vivo* by imaging *C. elegans* expressing fluorescent protein-tagged EV cargos. We discovered that loss of SOD-1, as well as the SOD-1(G85R) amyotrophic lateral sclerosis (ALS) pathogenic variant, increased EV shedding from the cilium distal tip, and this was associated with greater abundance of EV cargo in this ciliary compartment. In contrast, loss of SOD-1 reduced the glial uptake of a different cargo present in EVs shed from the ciliary base. Together, this suggests that redox balance has a subtype-specific effect on EV biogenesis, influencing neuron communication *in vivo*.

## 1. Introduction

Extracellular vesicles (EVs) are released by most, if not all, cells to mediate cell-to-cell transport of bioactive macromolecules. The protein, RNA, and metabolite cargo carried by EVs regulate physiological processes and the propagation of pathological conditions, including neurodegenerative diseases [1], [2], [3], [4]. Mutations in the Cu/Zn superoxide dismutase *SOD1* cause approximately 1 in 5 cases of familial amyotrophic lateral sclerosis (fALS), a fatal neurodegenerative disease characterized by loss of motor neurons [5], [6]. EVs derived from neurons and astrocytes that express pathogenic variants of *SOD1* are believed to contribute to the progression of ALS due to altered cargo content, including the presence of toxic aggregated SOD1 [7], [8]. In addition, EV biogenesis is enhanced in a swine model expressing pathogenic SOD1(G93A) and cultured astrocytes that overexpress this variant [9]. However, it is unclear whether this increase in EV release is due to the presence of toxic misfolded SOD1 or oxidative stress, as the majority of the *SOD1* ALS mutations also cause a reduction in enzymatic activity [10], [11].

SOD1 is an antioxidant enzyme that dismutates superoxide radicals (O2^•–^) into less harmful hydrogen peroxide (H_2_O_2_) and oxygen, such that SOD1 deficiency results in elevated O2^•–^, which generates oxidative stress within cells [12]. Children homozygous for loss-of-function *SOD1* mutations, as well as *Sod1* knockout mice have significant motor system deficits [13], [14], [15], [16], [17], [18], while loss of the *C. elegans* ortholog *sod-1* leads to degeneration of glutamatergic neurons upon exposure to cellular stress [10]. Pharmacological induction of oxidative stress can alter EV release in cell culture systems and redox-related proteins are cargo in EVs isolated from the biological fluids of many organisms [19], [20], [21], [22]. However, it is unknown whether SOD1 activity affects EV shedding and uptake. Since EV biogenesis occurs in response to signaling cues in native cellular environments, we sought to use an *in vivo* model to determine the impact of this antioxidant enzyme on EV release.

One site of EV shedding is the non-motile primary cilium, a specialized organelle protruding from non-dividing cells that serves as a platform for organizing both signal detection and transmission [23], [24]. Cilia are present on nearly all neurons in the vertebrate central and peripheral nervous systems, so it is critical to understand how oxidative stress regulates ciliary EV biogenesis [25], [26], [27]. In *C. elegans*, bioactive EVs termed ectosomes bud from the cilia of ray type B (RnB) sensory neurons in the *C. elegans* male tail and then are either taken up by the surrounding glia to mediate intercellular communication or discharged into the environment through a pore in the cuticle to facilitate animal-to-animal communication [24], [28], [29], [30], [31] (Fig. 1A). Using *C. elegans* that express fluorescently-tagged EV cargoes, we can quantify EV shedding from RnB cilia, the abundance of EV cargoes in the releasing cilia, and glial uptake of EV cargo. We discovered that SOD-1 is expressed in the RnB EV-releasing neurons (EVNs). Both the SOD-1(G85R) pathogenic mutation and a *sod-1* deletion mutation increased EV budding from the cilium distal tip. Loss of *sod-1* altered abundance of EV cargoes in the specific ciliary compartments where these proteins are packaged into EVs. Further, deletion of *sod-1* reduced the glial uptake of EVs derived from the ciliary base. Together, our results demonstrate that loss of SOD-1 function impacts EV shedding, cargo abundance, and uptake.

**Figure 1.**
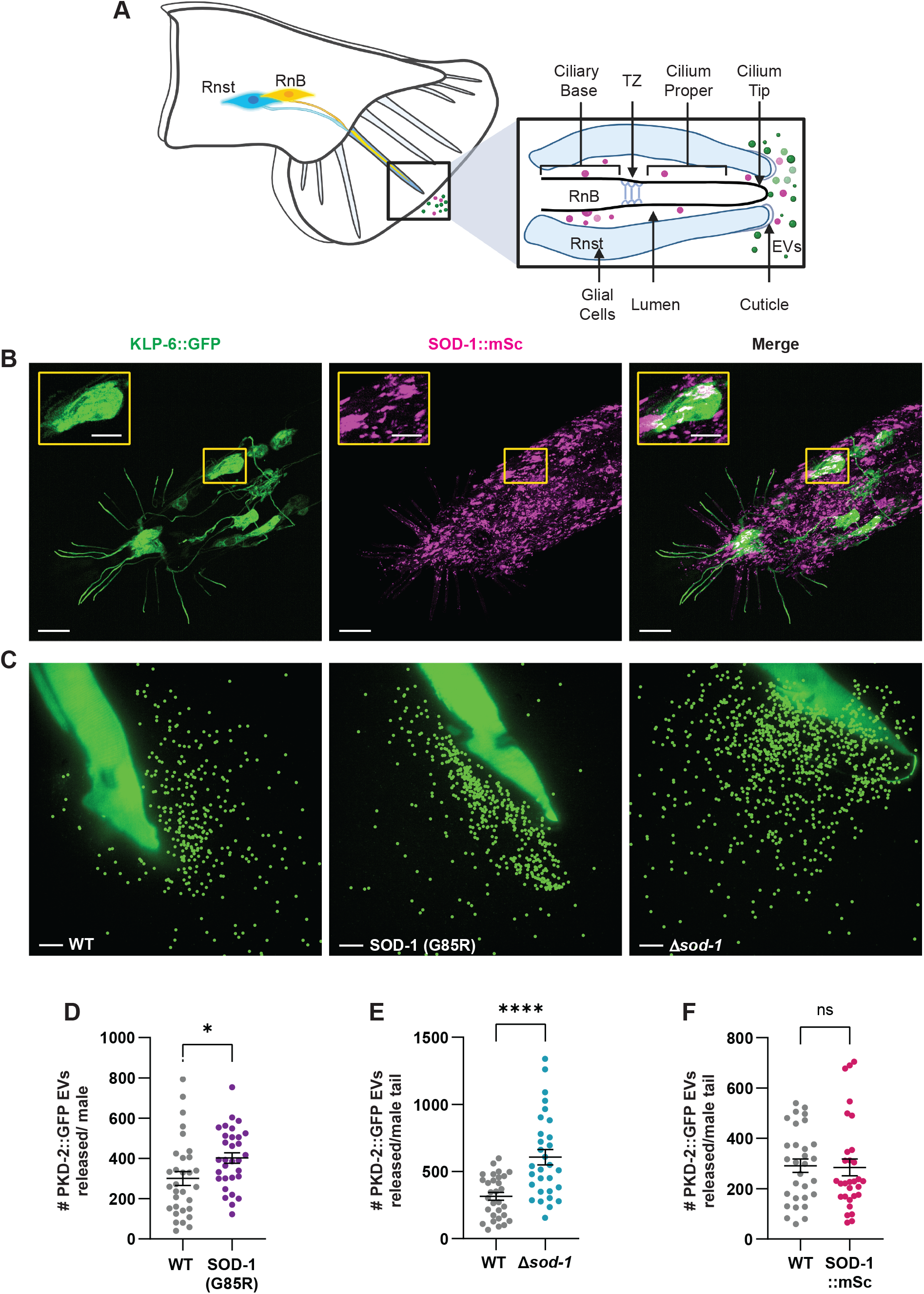
SOD-1 impacts release of PKD-2-containing EVs. (A) Schematic depicting EV shedding from the ciliary base (magenta EVs) and cilium distal tip (green EVs) of *C. elegans* male tail RnB sensory neurons, followed by release from the luminal space into the environment. (B) Endogenously tagged SOD-1::mSc coexpressed with KLP-6::GFP in RnB neurons of the male tail. Scale, 10 μm; inset scale, 5 μm. (C) Representative images of PKD-2::GFP (*henSi20*) EVs released from wild-type (left), SOD(G85R) (middle), and *sod-1(tm776)* deletion mutant (right) animals. EVs are marked with spots (Imaris software) for visualization; original, non-cropped, unmarked images are in Sup. Fig. S1; scale, 10 μm. (D) SOD-1(G85R) caused an increase in PKD-2::GFP EV shedding; n = 31. (E) Loss of *sod-1* increased PKD-2::GFP EV release in *sod-1(tm776)* animals; n = 30. (F) Addition of the mScarlet tag to SOD-1 did not affect PKD-2::GFP EV release. Data are presented as mean +/-SEM, Student’s t-test for (D,E), Mann-Whitney test (F), * p < 0.05, **** p<0.0001, ns indicates not significant.

## 2. Materials and Methods

### 2.1. *C. elegans* strains and maintenance

All strains were cultured at 20°C on Nematode Growth Media (NGM) plates seeded with OP50 *E. coli. sod-1(tm776)* II, *sod-1(hen25)* II, *sod-1(hen39)* [SOD-1(G85R)] and *him-5(e1490)* V are mutant alleles. *sod-1*(*syb8605* [*sod-1*::*mSc*]) II, *mks-2(oq101* [*mks-2*::*mNG*]) II, and *mks-2(syb7299* [*mks-2::mSc*]) II are endogenous insertion alleles. *myIs10* [*klp-6 promoter::klp-6::GFP*] and *nsls198* [*mir-228 promoter::GFP*] are integrated transgenes. *henSi17* [*klp-6 promoter::clhm-1::tdT*] III and *henSi20* [*pkd-2 promoter::pkd-2::GFP*] IV are single copy transgenes. All strains contain *him-5(e1490)*; control strains with no additional mutant allele present are referred to as wild type. See Supporting Information, Supplemental Table S1 for a list of all strains used in this work.

### 2.2. Cilia and glial imaging and analysis

24 hours post fourth larval (L4) stage, *C. elegans* males were immobilized with 50 mM levamisole (ThermoFisher Cat #: AC187870100) on 3% agarose pads on microscope slides. Z-stack images of splayed male tails were acquired with an Andor Dragonfly microscope (63x objective) and Zyla sCMOS camera. Identical image acquisition settings were used for all images that were directly compared; control and experimental strains were always imaged on the same day with the same settings.

Imaris (Oxford Instruments) was used for volumetric and fluorescence intensity analysis of cilia 3D reconstructions as described [32]. Briefly, ROIs were drawn around the ciliary region and preset intensity thresholds (fluorescent protein dependent) were applied to identify the ciliary base and cilium proper with the “Surface” tool. For ciliary length, a ROI was drawn to enclose KLP-6::GFP-labeled cilium proper, up to the edge of the MKS-2::mSc-labeled TZ. The length or cilia was traced and measured with “filament” function (segment seed threshold = 60). For measurement of CLHM-1::tdT engulfed by glial cells, surfaces were first generated to encapsulate the glial cells marked by GFP expression from the *mir-228* promoter in 3D reconstructions of Z-stack images. CLHM-1::tdT puncta were then identified using the Imaris “Surface” function. CLHM-1::tdT puncta that were localized within the ciliary region as well as those that did not colocalize with the glial GFP were manually excluded from the analysis.

### 2.3. EV imaging and analysis

Eight transgenic L4 hermaphrodites from control and mutant strains were picked onto 6 cm NGM plates and allowed to propagate for 4 days. Before imaging, adult males were briefly placed onto unseeded NGM plates to remove residual *E. coli* contamination, then immobilized in 20 mM levamisole (100 mM diluted in IMage-iT FX Signal Enhancer medium; ThermoFisher Item no. I36933) on 3% agarose pads on slides and covered with high-performance cover glass (Zeiss, Item no. 474030-9020-000). Slides and coverslips were pre-cleaned with ethanol, rinsed in HPLC water, and dried on heat block to prevent dust contamination [32]. EV images were collected with an Andor Dragonfly microscope and Andor Zyla sCMOS detector using total internal reflection fluorescence (TIRF) microscopy; the critical angle was optimized for each animal. Imaging was performed 40 ± 5 minutes after animal mounting, with control and experimental strains always imaged on the same day. EVs were quantified in Imaris software (Oxford Instruments) using the “Spot” function (object size, 0.350 μm diameter) and dataset-specific quality threshold determined by analysis of negative controls.

### 2.4. Male mating behaviors with freely moving hermaphrodites

L4 males were picked onto separate plates the day before mating assays. Mating plates were prepared by placing two 2.5 μl drops of freshly cultured *E. coli* OP50 on low peptone NGM plates. The next day, ten adult hermaphrodites were picked onto the mating plates and allowed to acclimate on the bacterial lawns. A single age-synchronized adult male worm was introduced onto the plate and recorded for 15 min. using an AmScope (MU1000) camera and Amlite software. Male behaviors in the recordings including ventral touch time (min), vulva touch time (min), and the number of successful and failed turns were analyzed as described [33].

### 2.5. Statistical Analysis

GraphPad Prism version 10 was used for statistical analysis and graphing. Depending on dataset normality, determined using the Anderson-Darling normality test, either the Student’s *t* test or Mann-Whitney U test was used (*p< 0.05, **p< 0.01, ***p< 0.001, ****p<0.0001). Figure legends specify statistical details for each experiment.

## 3. Results

### 3.1. Loss of SOD-1 function increases release of EVs from the cilium distal tip

Due to its important function as an antioxidant enzyme, *sod-1* is widely expressed in *C. elegans* [34]. To more clearly define the expression and localization pattern for SOD-1, we inserted a C-terminal mScarlet tag at the *sod-1* endogenous locus (SOD-1::mSc) using CRISPR/Cas9 genome editing. Co-expression of SOD-1::mSc with GFP-tagged KLP-6, which is expressed in the specialized EV-releasing neurons (EVNs) [28], [35], showed that SOD-1 puncta were present in male-specific RnB and HOB neurons as well as in many other cells throughout the body (Fig. 1B).

Having established that SOD-1 is expressed in the EVNs, we sought to determine if mutations in *sod-1* affect shedding of ciliary EVS, termed ectosomes. The TRP polycystin ion channel PKD-2 is a cargo in EVs that are primarily shed from the cilium distal tip [28], [30], [32]. We generated strains expressing GFP-tagged PKD-2 from a single copy transgene in a high incidence of males *him-5(e1490)* background, which causes elevated frequency of X chromosome non-disjunction, resulting in an increase in male progeny [36]. One strain contained the SOD-1(G85R) ALS-associated mutation [37], another strain included the *sod-1(tm776)* deletion mutation, and the control strain was wild type at the *sod-1* locus. Using total internal reflection fluorescence (TIRF) microscopy, we imaged EVs released into the environment from the male tail EVNs. We discovered that both the SOD-1(G85R) mutation and deletion of *sod-1* significantly increased the shedding of the cilium tip-derived PKD-2-containing EVs when compared to the control (Fig. 1C-E; Sup. Fig. S1). As these mutants both exhibited the same phenotype, this suggests that the impact of the ALS-associated mutation on EV shedding is due to loss of enzyme function.

To assess where SOD-1 could be acting to have this effect, we further analyzed the localization pattern of SOD-1::mSc. First, we determined that the SOD-1::mSc fusion protein is functional, as addition of the C-terminal mScarlet tag did not alter release of PKD-2-containing EVs into the environment (Fig. 1F). SOD-1::mSc puncta were observed in the EVN cell bodies and occasionally the dendrites, but were absent from the ciliary region (Fig. 1B). This suggests that SOD-1 does not act directly in the cilium to regulate EV shedding, but rather, an increase oxidative stress resulting from loss of superoxide dismutase activity likely heightens biogenesis of ciliary ectosomes.

### 3.2. SOD-1 affects the abundance of different EV cargoes in ciliary compartments

Shedding of PKD-2::GFP-labeled EVs is a dynamic process that can be influenced by the abundance of PKD-2 in the cilium [28], [30], [33]. To investigate whether the increase in PKD-2 EV release could be due to altered enrichment of the PKD-2 EV cargo in the *sod-1* mutant cilia, we quantified PKD-2::GFP fluorescence intensity and volume in the cilium proper and ciliary base. All strains also contained mScarlet-tagged MKS-2 (MKS-2::mSc), a protein in the Meckel syndrome (MKS) complex that localizes to the transition zone (TZ) [38], which enabled us to clearly distinguish the different ciliary compartments (Fig. 2A-B). Loss of *sod-1* had no effect on PKD-2 abundance in the ciliary base, but did increase enrichment of PKD-2::GFP in the cilium proper, which may underlie the increase in EV release from cilium distal tip observed in the *sod-1* mutants. Oxidative stress can also impact primary cilium length [39], [40]. To determine if the increase in PKD-2::GFP volume is due to a change in the length of the RnB cilia, we expressed KLP-6::GFP to fill out the EVNs and MKS-2::mSc to label the TZ, then measured cilia in *sod-1* deletion mutant and control animals (Fig. 2E). Loss of *sod-1* did not alter cilium length (Fig. 2F; Sup. Fig. S2) or disrupt the sensory function of the RnB cilia, as *sod-1* mutants did not exhibit defects in mating behaviors (Sup. Fig. S3). Thus, SOD-1 activity contributes to EV-mediated signaling without affecting cilium morphology.

**Figure 2.**
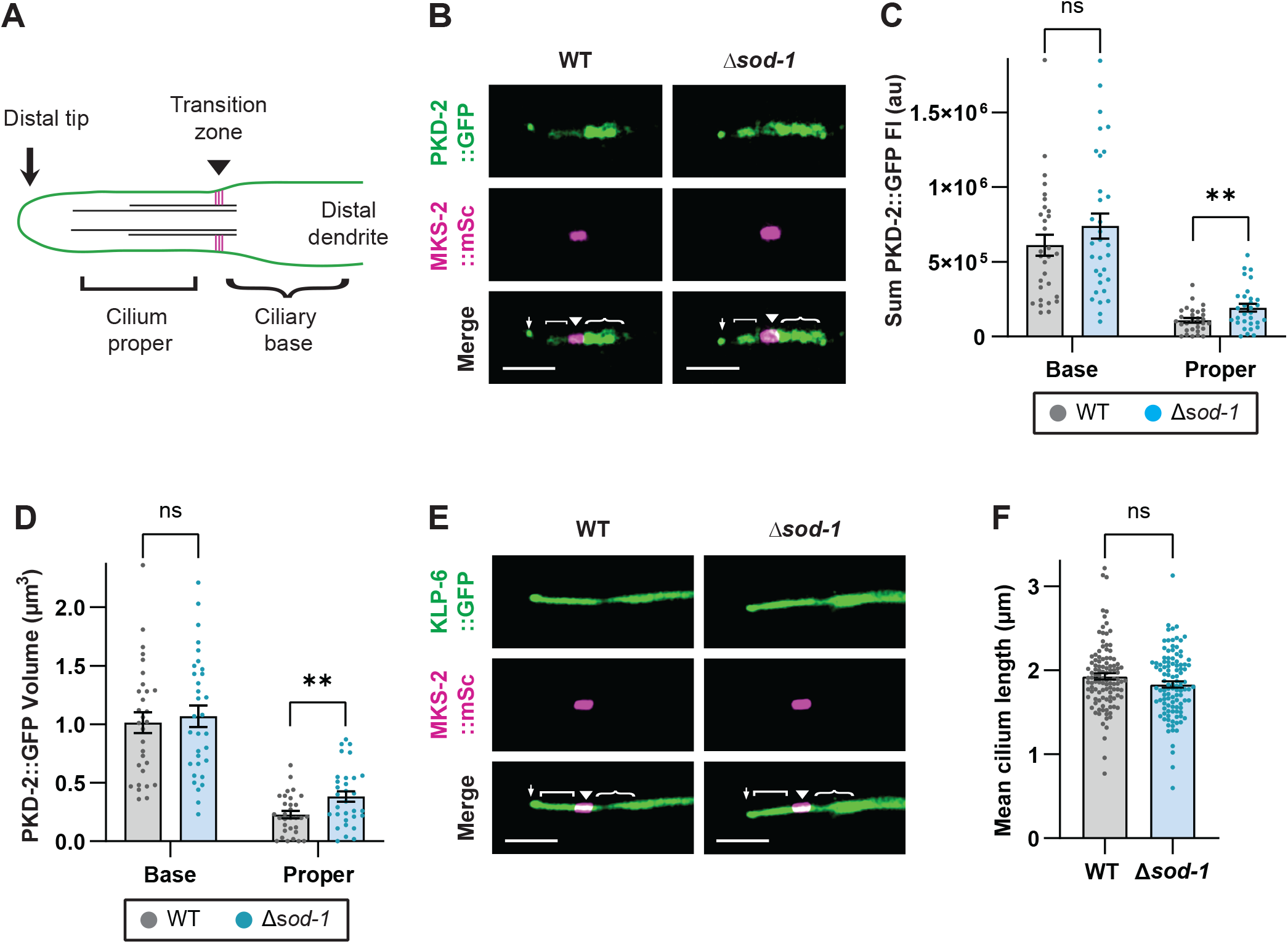
Loss of *sod-1* affects PKD-2 abundance in RnB cilia. (A) Schematic depicting ciliary subcompartments. The cilium distal tip (↓) oriented to the left, cilium proper ([), transition zone (▾), and ciliary base ({) are indicated here and in all images. (B) PKD-2::GFP (top) and TZ marker MKS-2::mSc (middle) in representative wild type (left) and *sod-1(hen25)* deletion mutant (right) cilia; scale, 2 μm. (C) Sum fluorescence intensity and (D) volume of PKD-2::GFP in the ciliary base and cilium proper in WT (grey) and *sod-1* mutant (blue) animals; n ≥ 30. (E) Representative images of R4B filled out with KLP-6::GFP (top) and the MKS-2::mSc labeled TZ (middle) in wild type (left) and *sod-1* mutant (right) animals; scale, 2 μm. (F) Cilium length was unchanged in the *sod-1* mutant compared to the wild type; n = 109. In this figure, data are shown as mean +/-SEM; Mann-Whitney test (C and F), Student’s t-test (D), ** p< 0.01.

Using *C. elegans* that express different fluorescent protein-tagged EV cargoes, we have shown that multiple distinct EV subpopulations are shed from the RnB neurons into the environment [28]. We next sought to establish whether loss of *sod-1* disrupted the ciliary abundance of the calcium homeostasis modulator (CLHM-1) ion channel, which is a cargo in EVs shed from the ciliary base [28]. Analysis of tdTomato-tagged CLHM-1 (CLHM-1::tdT) in the RnB neurons showed that loss of *sod-1* significantly reduced CLHM-1::tdT volume and intensity in the ciliary base, but had no impact on CLHM-1 abundance in the cilium proper (Fig. 3A-C). We next quantified the number of CLHM-1 EVs released into the environment from SOD-1(G85R) and *sod-1* deletion mutants and found no significant difference when compared to the control (Fig. 3D-F). Thus, despite reducing ciliary abundance of CLHM-1 in the ciliary base, loss of *sod-1* does not affect the shedding of CLHM-1 EVs from this ciliary compartment into the environment.

**Figure 3.**
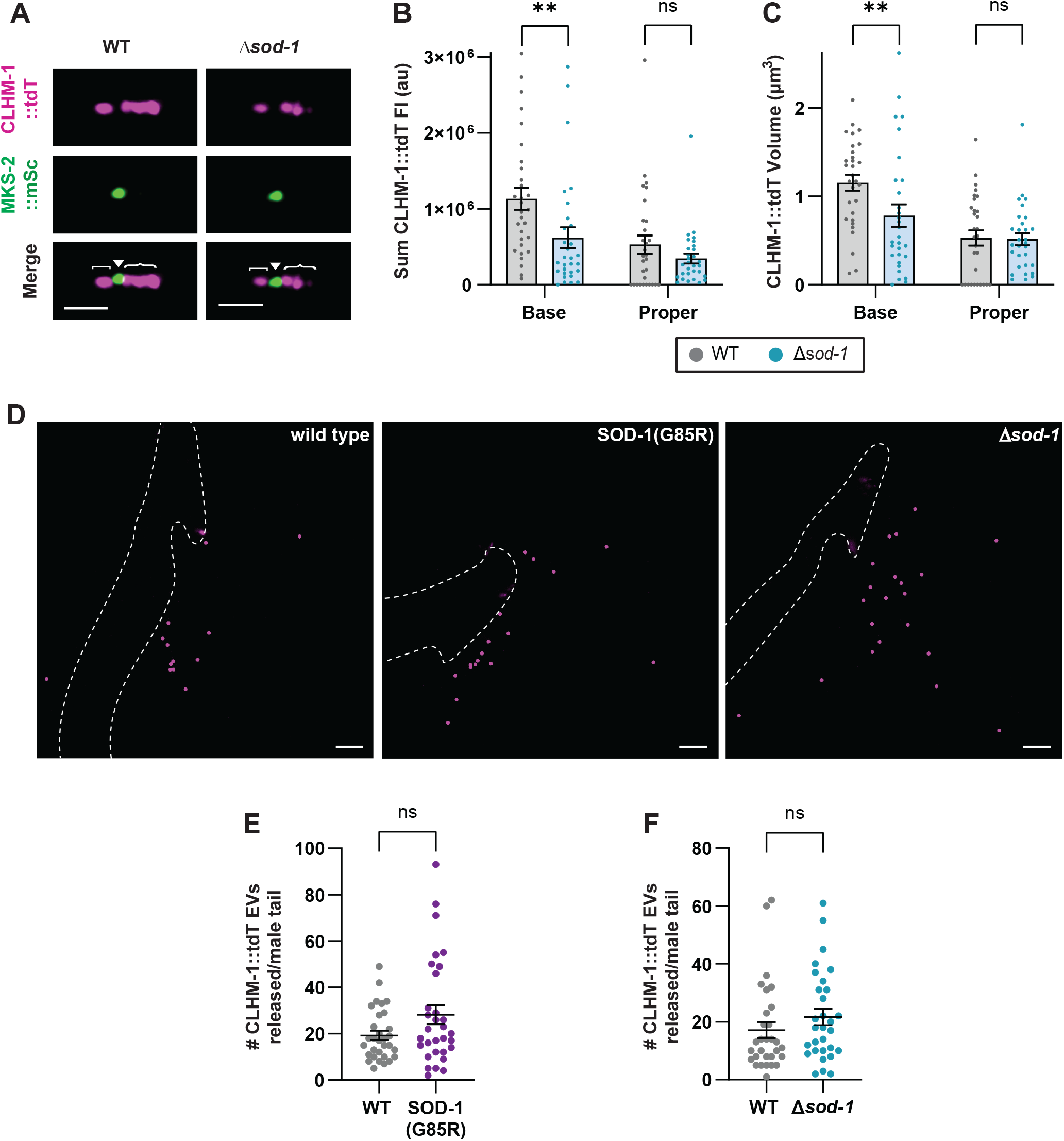
Loss of *sod-1* reduces CLHM-1 abundance in the ciliary base, but does not impact EV shedding. (A) Representative images of CLHM-1::tdT (top) and MKS-2::mNG (middle) in wild-type and *sod-1(hen25)* mutant R5B cilia. Cilium proper ([), transition zone (▾), ciliary base ({) and scale, 2 μm, are indicated. (B) CLHM-1::tdT sum fluorescence intensity and (C) volume in the ciliary base and cilium proper of wild-type (grey) and *sod-1* mutant (blue) cilia; n ≥ 29. (D) Representative images of CLHM-1::tdT (*henSi17*) EVs shed from wild type, SOD-1(G85R) and *sod-1(hen25)* mutant male tails; scale, 10 μm. EVs marked with spots (Imaris software) for visualization; see also Sup. Fig. 1. (E) SOD-1(G85R) and (F) deletion of *sod-1* does not impact CLHM-1::tdT EV release into the environment; n ≥ 30. Data are mean ± SEM; Mann-Whitney test, ** p< 0.01.

#### SOD-1 activity affects glial uptake of ciliary base-derived EVs

EVs shed from the ciliary base of sensory neurons in the *C. elegans* head can be phagocytosed by surrounding glial cells [29]. To explore whether CLHM-1-containing EVs can be taken up by Rnst support cells, we created a strain with CLHM-1::tdT expressed only in the EVNs and GFP expressed from the *mir-228* promoter in the Rnst glia [41]. CLHM-1::tdT was present in 3D reconstructions of the GFP positive glial cells, demonstrating that Rnst support cells engulf CLHM-1 EVs (Fig. 4A-C). Next, we investigated whether loss of *sod-1* affected the abundance of the CLHM-1::tdT EV cargo in the Rnst glia. CLHM-1::tdT volume, puncta number, and sum intensity in the Rnst support cells were significantly reduced in the *sod-1* mutants compared to wild type (Fig. 4D-F). Together, these results suggest that while loss of *sod-1* does not affect release of CLHM-1 EVs into the environment, it leads to reduced abundance of this EV cargo in the surrounding glia.

**Figure 4.**
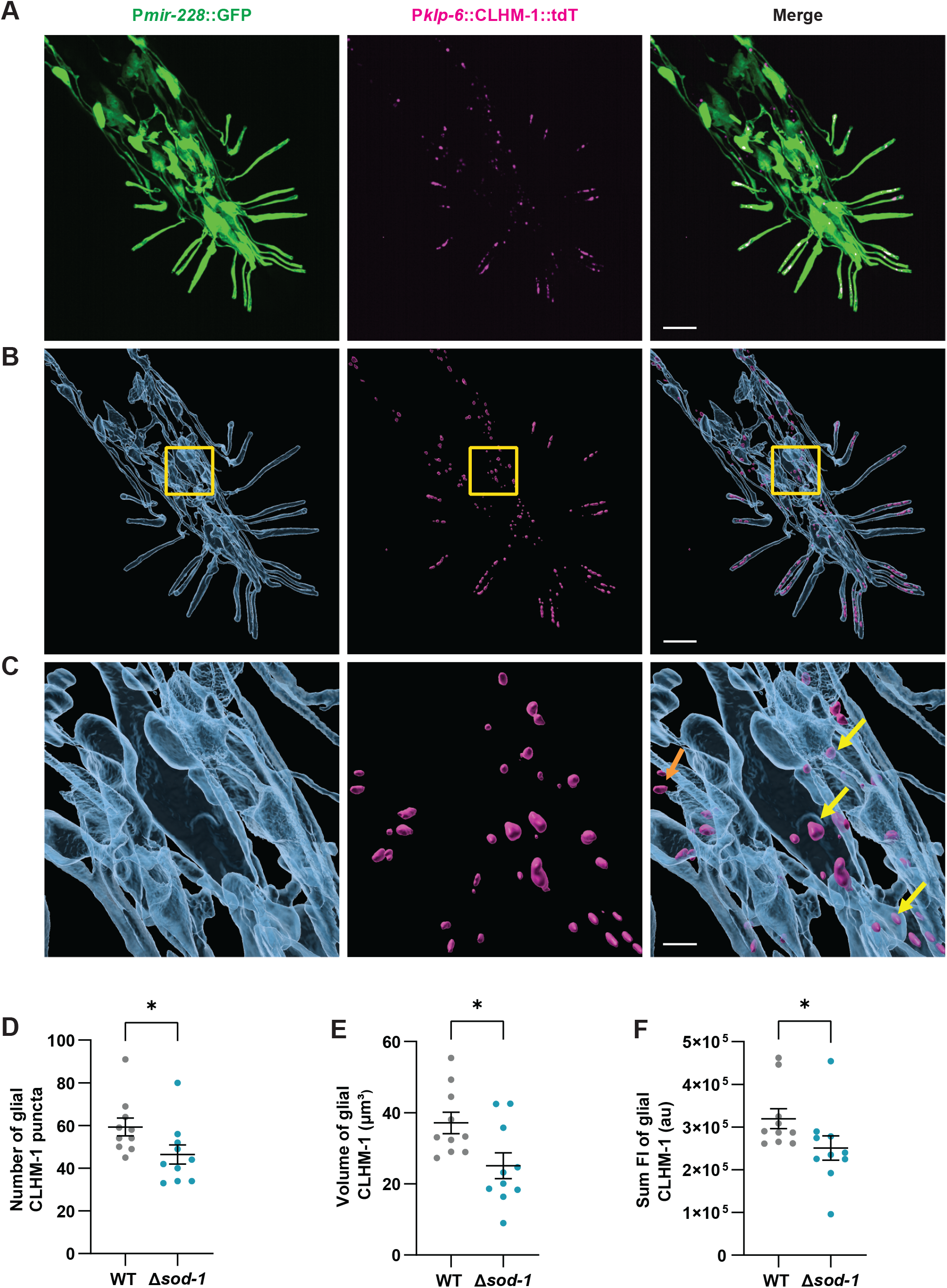
Glial uptake of CLHM-1 EVs is altered by the loss of *sod-1*. (A) Representative image of GFP expressed in Rnst glia from the *mir-228* promoter (left) and CLHM-1::tdT expressed in the RnB and HOB neurons from the *klp-6* promoter (middle); scale, 10 µm (B) Image from A, with Rnst glia and CLHM-1::tdT labeled with surfaces for visualization. (C) Corresponding to the boxed region in B, CLHM-1::tdT puncta are observed inside the opaque surface defining the Rnst (yellow arrows) as well as outside of the glia (orange arrow), in the RnB cell bodies; scale, 2 µm. Total number of glial CLHM-1::tdT (D) puncta, (E) volume, and (F) average sum intensity in wild-type and *sod-1(tm776)* mutant animals; n = 10. All data are presented as mean ± SEM, Student’s t-test (D,E), Mann-Whitney test (F) * p < 0.05.

## 4. Discussion

Pathogenic mutations in the antioxidant enzyme SOD1 lead to enrichment of redox-related cargos and pathogenic misfolded proteins in EVs [7], [8]; however, the impact on EV release and uptake *in vivo* has not been established. Using fluorescently tagged cargos to visualize EV shedding from *C. elegans* sensory neurons, we discovered that loss of *sod-1* increased biogenesis of PKD-2-containing ectosomes from the cilium distal tip, without affecting cilium length or sensory function. Intriguingly, *sod-1* mutations had a different impact on the CLHM-1 EV cargo, reducing the abundance of this protein in the ciliary base and the amount of CLHM-1 taken up by surrounding glia. Since the environmental release of the EVs containing CLHM-1 was unaffected, this suggests that loss of *sod-1* specifically reduces the EV-mediated transfer of this cargo to the glia. Together, our findings demonstrate that loss of SOD-1 function affects EV-mediated signaling by exerting distinct effects on different EV subpopulations.

While SOD-1 was previously shown to localize to the sensory cilia of the ASE neurons in the *C. elegans* head [34], SOD-1 was not present in the RnB cilia. Thus, the increase in shedding of EVs from the cilium distal tip in *sod-1* mutants may instead result from disruption of redox balance. Oxidative stress can have a complex effect on ciliogeneisis and cilium length, with the specific outcome dependent on ROS level [39], [40], [42]. However, the RnB cilia in the *sod-1* mutant do not exhibit altered length or sensory function, suggesting that the elevated EV shedding from the distal tip does not result from defects in cilium morphology. Further supporting this, mutants with altered microtubule isotypes and post-translational modifications that disrupt the RnB cilia exhibit decreased EV release into the environment [43], [44]. While cilia are microtubule-based organelles, the ciliary actin network regulates ectocytosis from cilium distal tip and intraflagellar transport (IFT), which controls cargo entry and exit from the cilium [45], [46], [47], [48]. Since oxidative stress can damage the actin cytoskeleton [49], [50], we posit this as a possible mechanism by which loss of *sod-1* impacts distal tip EV shedding and ciliary cargo abundance. Alternatively, SOD1-dependent changes in gene expression and the proteome in response to oxidative stress could affect ectosome biogenesis [51], [52].

EV biogenesis from the ciliary base compared to the cilium distal dip is differentially regulated [28], [32]. Loss of *sod-1* reduced glial uptake of the CLHM-1, which is a cargo in EVs shed from the ciliary base. While it remains possible that this results from reduced CLHM-1 abundance in the periciliary membrane compartment, the release of ciliary base-derived EVs into the environment was unaffected in the *sod-1* mutants. Instead, oxidative stress in the receiving glial cells could impact EV targeting. In the future, cell specific depletion of *sod-1* in distinct cell types *in vivo* could be used to distinguish where SOD-1 acts to control neuron-glia communication.

Misfolded SOD1 aggregates in EVs contribute to spread of ALS pathogenesis in recipient cells [7], [8]. Here we show that in addition to the established gain of function effects, SOD-1(G85R) pathogenic mutations that cause loss of superoxide dismutase enzymatic function have a more broad impact on intercellular communication by altering EV shedding. Given that loss of SOD1 enzymatic activity causes oxidative stress, this suggests that more broadly, disruption of redox balance may be a key regulator of EV-mediated communication that could contribute to progression of neurodegenerative disease.

## Abbreviations

EV: SOD-1
TIRF: CLHM-1
PKD-2: EVN

## CRediT authorship contribution statement

**Nahin Siara Prova:** Conceptualization, Investigation, Formal analysis, Methodology, Data curation, Writing – original draft, Writing – review & editing. **Malek Elsayyid:** Investigation, Methodology. **Jessica E. Tanis:** Conceptualization, Writing – original draft, Writing – review & editing, Supervision, Funding acquisition.

## Acknowledgments

We thank the *Caenorhabditis* Genetics Center, which is supported by the NIH-ORIP (P40 OD010440), for strains and the University of Delaware BioImaging facility staff for their support. Microscopy access was supported by National Institutes of Health NIGMS P20 GM103446, NIH-NIGMS P20 GM139760, and the State of Delaware; equipment was acquired with an NIGMS S10 OD030321. This work was supported by the National Institutes of Health NIGMS R01 GM135433 (to J.E.T.).

## Supplemental Figure Legends

**Supplemental Figure S1.**SOD-1 impacts EV shedding from male tail EVNs. Representative images of (A) PKD-2::GFP *(henSi20)* and (B) CLHM-1::tdT *(henSi17)* labeled EVs released from wild type (left), SOD-1 G85R (middle), and *sod-1(tm776)* deletion mutant males. Images in A correspond to Fig. 1C and images in B correspond to Fig. 3D; scale, 10 µm.

**Supplemental Figure S2.**.Loss of *sod-1* does not impact cilium length (A) There was no significant difference in the length of the R3B (marginal opening), R4B (ventral opening), and R5B (dorsal opening) cilia in the *sod-1(tm776)* mutant compared to the wild-type animals; n ≥ 21 animals. Data are represented as mean +/-SEM: Mann-Whitney test.

**Supplemental Figure S3.**.Male mating behavior is not impacted by the loss of *sod-1* (A) Total turn attempts, (B) number of failed turns per minute of hermaphrodite contact time, and (C) the percent of successfully completed turns were unchanged in *sod-1(tm776)* compared to wild type. (D) Male stopping at hermaphrodite vulva and (E) ventral contact response was unaffected in *sod-1(tm776)* mutant animals. n ≥ 11, data are presented as mean ± SEM; Student’s t-test (A, D), Mann-Whitney test (B, C and E).

