## Supplemental Figures for "Superoxide dismutase impacts extracellular vesicle biogenesis and uptake"

### Supplemental Figure 1

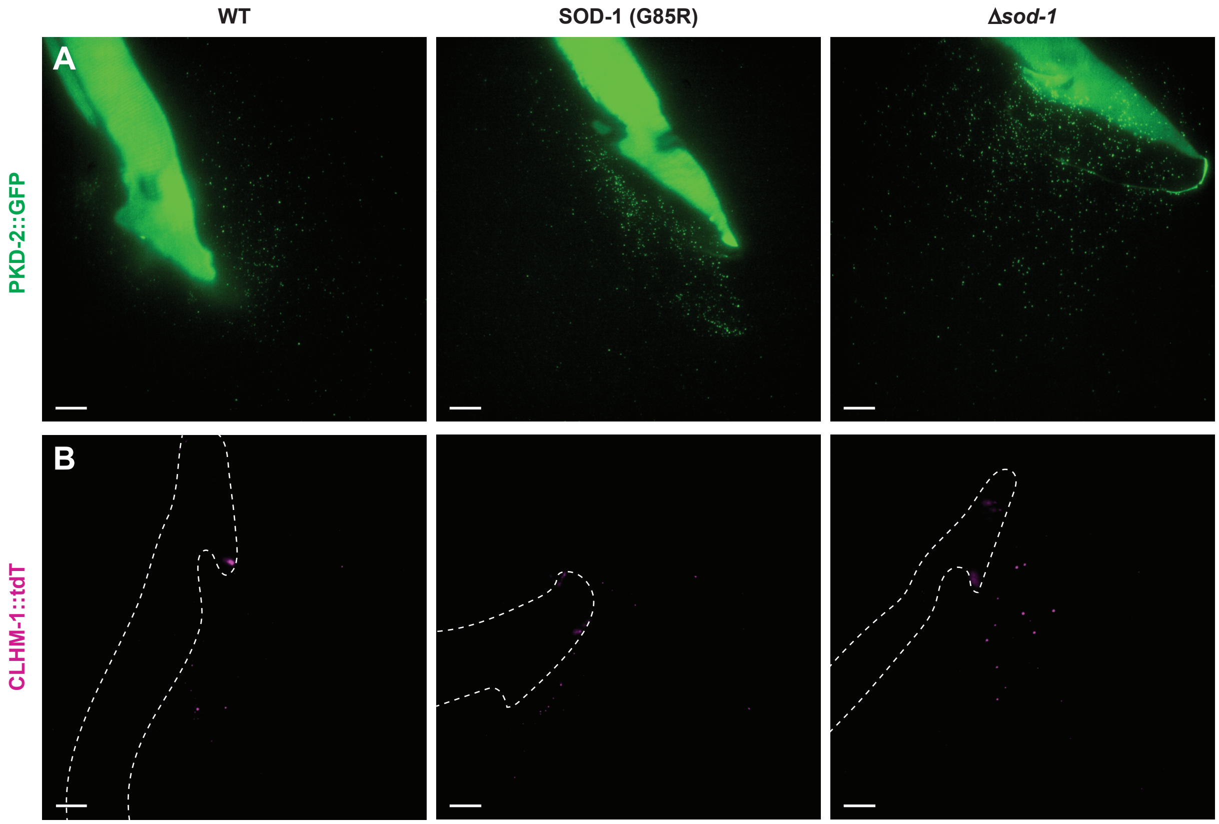


**Supplemental Figure S1. SOD-1 impacts EV shedding from male tail EVNs.** Representative images of (A) PKD-2::GFP (*henSi20*) and (B) CLHM-1::tdT (*henSi17*) labeled EVs released from wild type (left), SOD-1 G85R (middle), and *sod-1(tm776)* deletion mutant males. Images in A correspond to Fig. 1C and images in B correspond to Fig. 3D; scale, 10 μm.

### Supplemental Figure 2

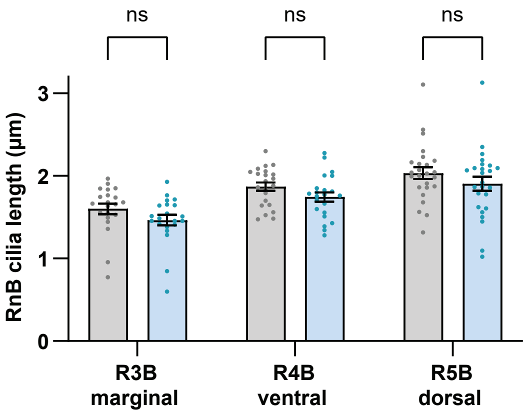


**Supplemental Figure S2. Loss of sod-1 does not impact cilium length** (A) There was no significant difference in the length of the R3B (marginal opening), R4B (ventral opening), and R5B (dorsal opening) cilia in the *sod-1(tm776)* mutant compared to the wild-type animals; n ≥ 21 animals. Data are represented as mean +/- SEM: Mann-Whitney test.

### Supplemental Figure 3

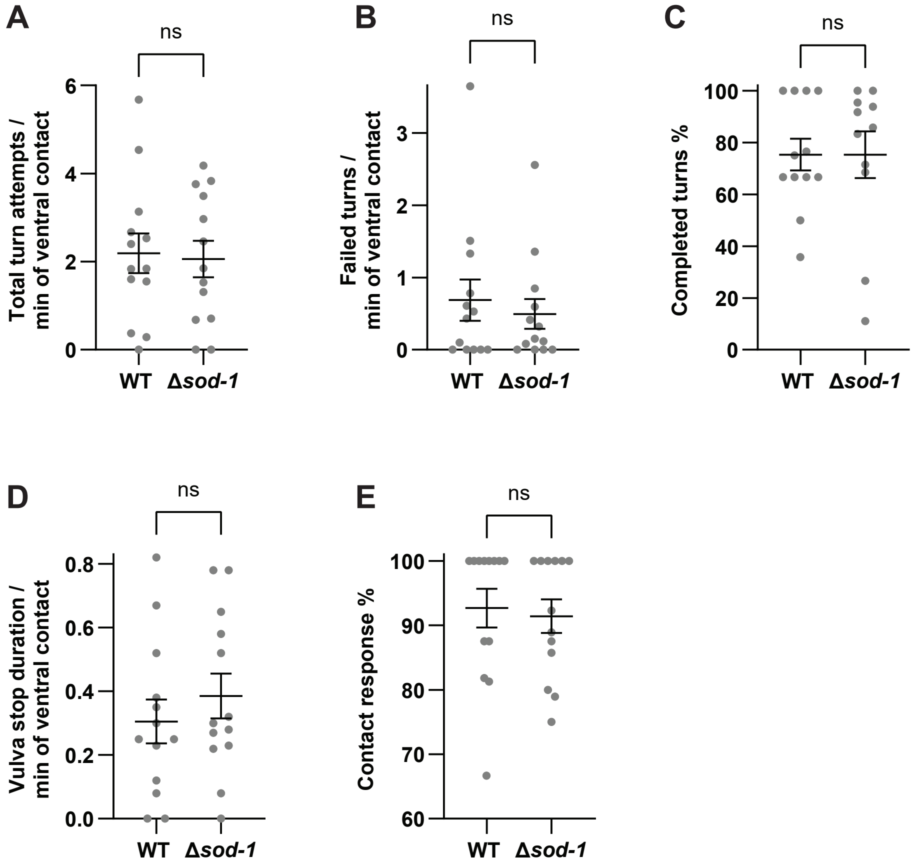


**Supplemental Figure S3. Male mating behavior is not impacted by the loss of *sod-1*** (A) Total turn attempts, (B) number of failed turns per minute of hermaphrodite contact time, and (C) the percent of successfully completed turns were unchanged in *sod-1(tm776)* compared to wild type. (D) Male stopping at hermaphrodite vulva and (E) ventral contact response was unaffected in *sod-1(tm776)* mutant animals. n ≥ 11, data are presented as mean ± SEM; Student's t-test (A, D), Mann-Whitney test (B, C and E).
